# *Salmonella enterica* genomes recovered from victims of a major 16th century epidemic in Mexico

**DOI:** 10.1101/106740

**Authors:** Åshild J. Vågene, Michael G. Campana, Nelly M. Robles García, Christina Warinner, Maria A. Spyrou, Aida Andrades Valtueña, Daniel Huson, Noreen Tuross, Alexander Herbig, Kirsten I. Bos, Johannes Krause

## Abstract

Indigenous populations of the Americas experienced high mortality rates during the early contact period as a result of infectious diseases, many of which were introduced by Europeans. Most of the pathogenic agents that caused these outbreaks remain unknown. Using a metagenomic tool called MALT to search for traces of ancient pathogen DNA, we were able to identify *Salmonella enterica* in individuals buried in an early contact era epidemic cemetery at Teposcolula-Yucundaa, Oaxaca in southern Mexico. This cemetery is linked to the 1545-1550 CE epidemic locally known as “*cocoliztli*”, the cause of which has been debated for over a century. Here we present two reconstructed ancient genomes for *Salmonella enterica* subsp. *enterica* serovar Paratyphi C, a bacterial cause of enteric fever. We propose that *S.* Paratyphi C contributed to the population decline during the 1545 *cocoliztli* outbreak in Mexico.

**One Sentence Summary:** Genomic evidence of enteric fever identified in an indigenous population from early contact period Mexico.

## Main Text

A variety of plants, animals, cultures, technologies and infectious diseases accompanied the movement of people from the Old World to the New World immediately following initial contact, in a process commonly known as the “Columbian exchange” (*1*). Old World infectious diseases introduced to the New World led to successive outbreaks that continued well into the 19^th^ century, causing high mortality and demographic collapse amongst indigenous populations (*2-6*). Rates of population decline linked to regionally specific epidemics are estimated to have reached as high as 95% (*4*), and the genetic impact of these diseases based on recent population-based studies of ancient and modern human exome and mitochondrial data attest to their scale (*7, 8*). One hypothesis assumes that the increased susceptibility of New World populations to Old World diseases facilitated European conquest, whereby rapidly disseminating diseases severely weakened indigenous populations (*3*), in some cases even in advance of European presence in the region (*3, 9*). Well-characterized Old World diseases such as smallpox, measles, mumps, and influenza have been accepted as the causes of later contact era outbreaks; however, the diseases responsible for many early contact period New World epidemics remain unknown, and have been the subject of scientific debate for over a century (*4-6, 9-12*).

Morphological changes in skeletal remains (*13*) and ethnohistorical accounts (*14*) are often explored as sources for understanding population health in the past, although both provide only limited resolution and have generated speculative and at times conflicting hypotheses about the diseases introduced to New World populations (*3, 4, 10, 12, 15, 16*). Most infectious diseases do not leave characteristic markers on the skeleton due to either their short periods of infectivity, the death of the victim in the acute phase before skeletal changes formed, or a lack of osteological involvement (*17*). While historical descriptions of infectious disease symptoms can be very detailed, they are subject to cultural biases, suffer from inaccuracies in translation, lack a foundational knowledge of germ theory, and describe historical forms of a condition that may differ from modern manifestations (*11, 15*).

Genome-wide studies of ancient pathogens have proven instrumental in both identifying and characterizing past human infectious diseases. For the most part, these studies have been restricted to skeletal collections where individuals display physical changes consistent with particular pathogens (*18-20*), where an historical context links a specific pathogen to a known epidemiological event (*21*), or where an organism was identified via targeted molecular screening without prior indication of its presence (*22*). Recent attempts to circumvent these limitations have concentrated on broad-spectrum molecular screening approaches focused on pathogen detection via fluorescence-hybridization-based microarray technology (*23*), identification via DNA-enrichment of certain microbial regions (*24*) or computational screening of non-enriched sequence data against human microbiome data sets (*25*). While these approaches offer improvements, they remain biased in the bacterial taxa used for species-level assignments. As archeological material is expected to harbor an abundance of bacteria that stem from the depositional context, omission of environmental taxa in species assignments can lead to falsepositive identifications.

Here we demonstrate the utility of a novel and comprehensive ancient pathogen screening approach that analyzes non-enriched short read DNA sequence data via high-throughput taxonomic assignment. The Megan ALignment Tool (MALT) is a rapid computational application used to generate metagenomic profiles by assigning sequenced reads in a dataset to their taxonomic nodes of best fit within a specified database (*26*). Here we use a curated reference database of 2783 bacterial genomes, comprising the entirety of those available in NCBI RefSeq (December 2015). Our approach limits ascertainment biases and false positive assignments that could result from databases deficient in environmental taxa. We applied MALT to non-enriched DNA sequence data from the pulp chamber of teeth collected from indigenous individuals excavated at the site of Teposcolula-Yucundaa, located in the highland Mixteca Alta region of Oaxaca, Mexico (*27, 28*). The site contains both pre-contact and contact era burials, including the earliest identified contact era epidemic burial ground in Mexico (*28, 29*) (Fig. 1). The site is historically linked to the *cocoliztli* epidemic of 1545-1550, described as one of the principal epidemiological events responsible for the cataclysmic population decline of 16^th^ century Mesoamerica (*12, 28, 30*). This outbreak affected large areas of central Mexico and Guatemala, spreading perhaps as far south as Peru (*9, 28, 30*). Via the MALT screening approach, we were able to identify ancient *Salmonella enterica* DNA in the sequence data generated from this archaeological material, to the exclusion of DNA stemming from the complex environmental background. While the pathogenic cause of the *cocoliztli* epidemic is ambiguous based on ethnohistorical evidence (*10, 12, 28, 30*), we report the first molecular evidence of microbial infection with the reconstruction of two ancient *S. enterica* subsp. *enterica* serovar Paratyphi C (enteric fever) genomes isolated from epidemic-associated contact era burials.

**Figure 1.**
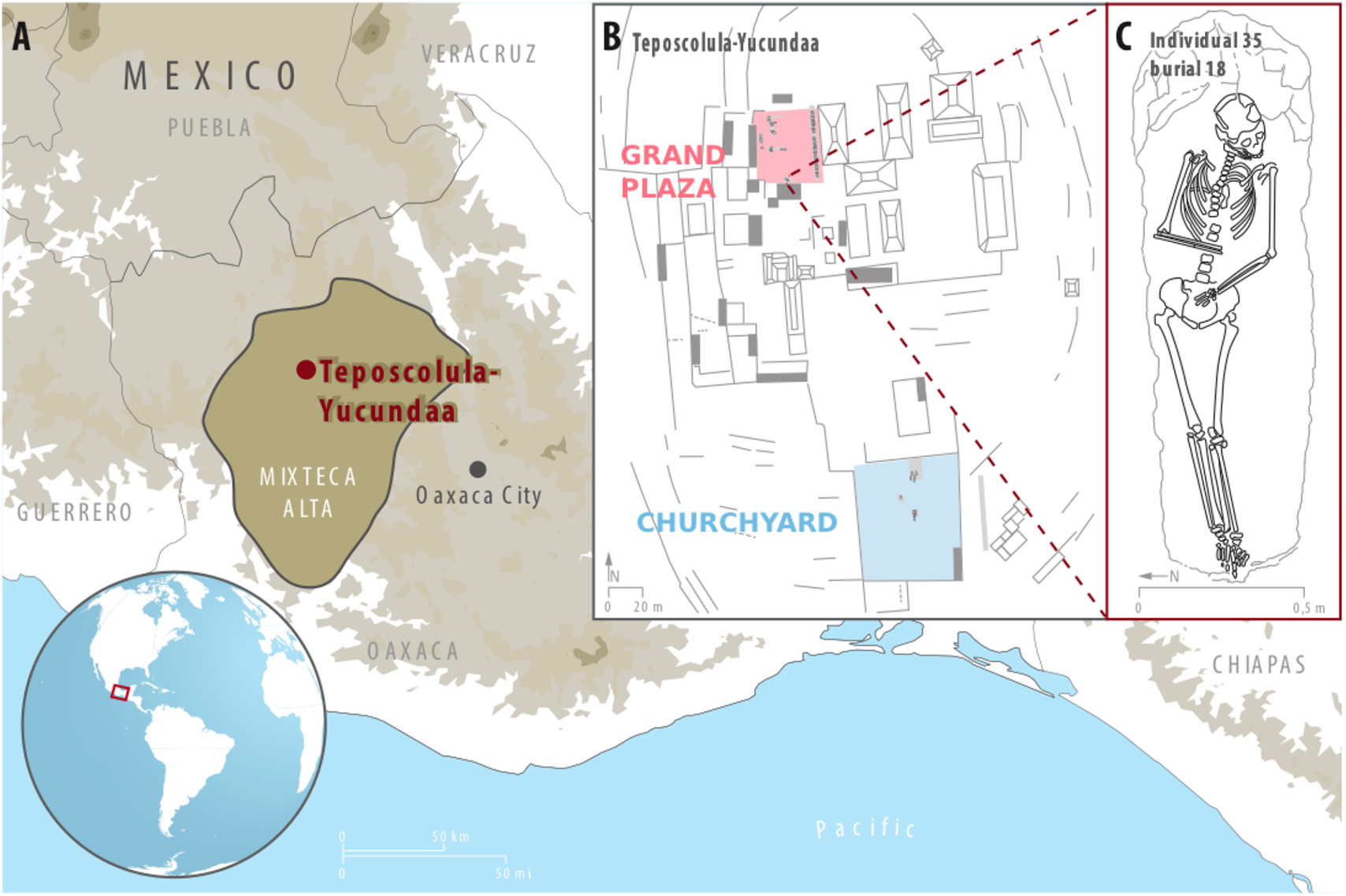
Overview of Teposcolula-Yucundaa. A) Illustrates the location of the Teposcolula-Yucundaa site in the Mixteca Alta region of Oaxaca, Mexico; B) central administrative area of Teposcolula-Yucundaa showing the relative positioning of the Grand Plaza and churchyard cemetery sites. Burials within each cemetery are indicated with dark grey outlines and the excavation area is shaded in grey; C) drawing of individual 35 from which the Tepos_35 *S.* Paratyphi C genome was isolated. Panels B and C were adapted from drawings provided by the Teposcolula-Yucundaa archaeological project archives-INAH and from (*28*).

The individuals included in this investigation were excavated from the contact era epidemic cemetery located in the Grand Plaza (administrative square) (n=24) and the pre-contact churchyard cemetery (n=5) at Teposcolula-Yucundaa between 2004 and 2010 (*28*) (Fig. 1; table S1; Supplementary materials S1). Previous work demonstrated ancient DNA preservation at the site through the identification of New World mitochondrial haplogroups in 48 individuals, 28 of which overlap with this study (*28*). Additionally, oxygen isotope analysis identified them as local inhabitants (*28*). Radiocarbon dating of selected individuals from both burial sites support archaeological evidence that the Grand Plaza and churchyard contain contact and pre-contact era burials, respectively (*29*). The Grand Plaza cemetery is estimated to contain more than 800 individuals, mostly interred in graves containing multiple individuals. Those that have been excavated exhibit a demographic profile consistent with an epidemic event (*27, 28*).

Tooth samples were processed according to protocols designed for ancient DNA work (Supplementary materials S2). An aggregate soil sample from the two burial grounds was analyzed in parallel to gain an overview of environmental bacteria that may have infiltrated our samples. Pre-processed sequencing data of approximately one million paired-end reads per tooth sample were analyzed with MALT (*26*) using a comprehensive database of all bacterial genomes available through NCBI RefSeq (December 2015) (Supplementary materials S3). MALT results were visualized in MEGAN (*31*) and taxonomic assignments were evaluated with attention to known pathogenic species. The total number of reads taxonomically assigned by MALT ranged from 3,694 to 33,586. Assigned reads belonging to bacterial constituents of human oral and soil microbiota are present in varying proportions amongst samples (Fig. 2; table S2). In three samples (Tepos_10, Tepos_14 and Tepos_35) 374 to 653 reads were assigned to *Salmonella enterica*, and of the *S. enterica* strains present in the database, *S*. Paratyphi C had the highest number of assigned reads. (Fig. 2; Supplementary materials S3; table S2). Mapping of these three samples to the *S.* Paratyphi C RKS4594 genome (NC_012125.1) revealed the characteristic pattern of damage expected of ancient DNA molecules (fig. S1; Supplementary materials S3; table S3). Subsequently, the sequencing data for all samples was mapped to the human genome (hg19), revealing a similar level of damage in the human reads for Tepos_10, Tepos_14 and Tepos_35, thus providing further support for the ancient endogenous origin of the *S. enterica* reads (Supplementary materials S7; table S7). An additional five individuals from the Grand Plaza cemetery and one negative control harbored low numbers of reads assigned to *S. enterica* ranging from 5 to 51 (table S2). These read counts were too low for further analysis and the five individuals were not included in downstream experiments. The negative control, however, continued to be included. No *S. enterica* reads were identified in any of the pre-contact churchyard samples, the soil sample, or the remaining negative controls (Fig. 2; Supplementary materials S3; table S2).

**Figure 2.**
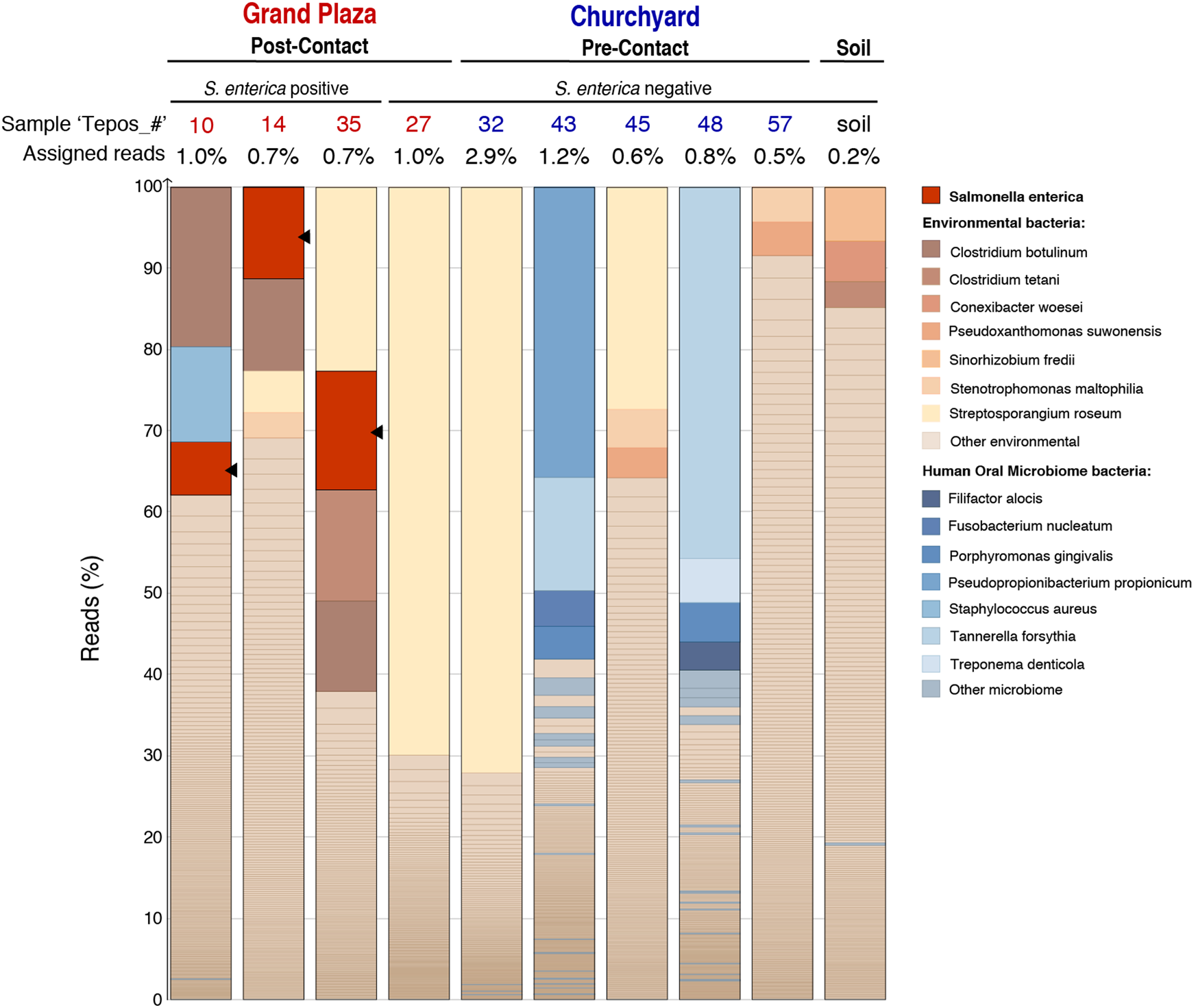
MALT analysis and pathogen screening of shotgun data. Shotgun data was analyzed with MALT (*26*) using a database constructed from all bacterial genomes available through NCBI RefSeq (December 2015). MALT results were visualized using MEGAN6 (*31*). The bar chart was constructed from the MEGAN6 output and is based on the percent reads assigned to bacterial species when using a 95% identity filter. Reads assigned to *Salmonella enterica* are colored red regardless. Other taxa to which 3% or more reads, per sample, were assigned are color-coded depending on whether they are ‘environmental’ or ‘human oral microbiome’ bacteria. Remaining taxa are sorted into two categories: ‘other environmental’ or ‘other microbiome’ (Supplementary materials S3). Samples from the post-contact Grand Plaza epidemic cemetery containing *S. enterica* reads, pre-contact samples from the churchyard cemetery and the soil sample are illustrated. Additionally, a sample negative for *S. enterica* from the Grand Plaza cemetery (Tepos_27) is shown. Samples whose names are colored in red are from the Grand Plaza and those in blue from the churchyard. The percentage of reads in the shotgun data assigned by MALT per sample is indicated at the top of each column.

**Figure 3.**
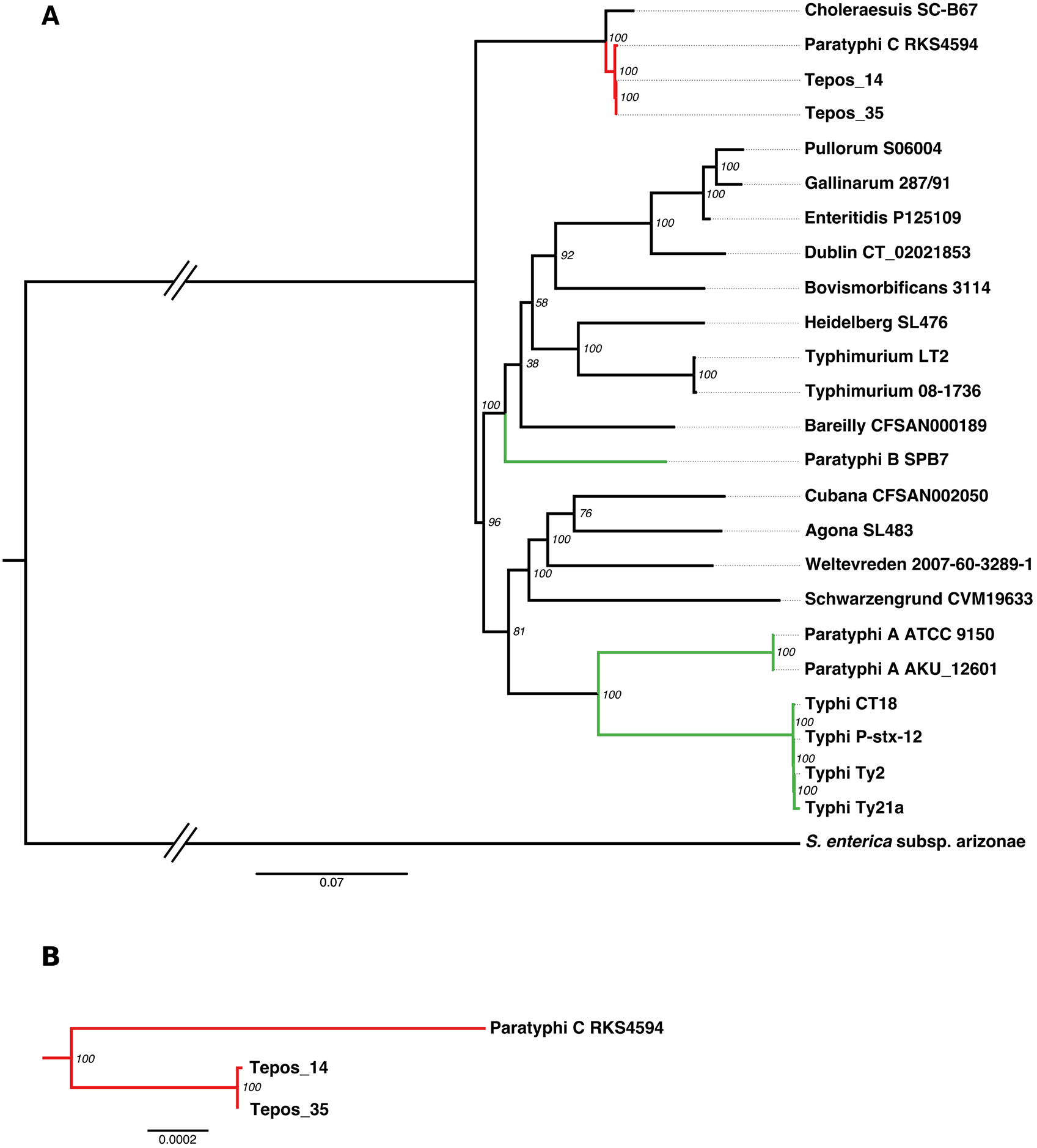
Maximum Likelihood *S. enterica* phylogeny. A) Maximum likelihood tree, based on the Tamura-Nei model, consisting of two ancient and 23 modern genomes. The tree is based on 81135 variant positions; positions with missing data were excluded. The two ancient and one modern *S.* Paratyphi C genomes are indicated in red. Other human specific strains that cause enteric fever, *S.* Typhi, *S.* Paratyphi A and *S.* Paratyphi B, are indicated in green. The nodes are labeled with bootstrap values. (B) An enlarged version of the *S.* Paratyphi C clade, illustrating the branch shortening of the two ancient genomes (Tepos_14 and Tepos_35).

To further authenticate and elucidate our findings we performed a whole-genome targeted array hybridization capture (*32, 33*), using probes designed to encompass modern *S. enterica* genome and plasmid diversity (table S4; Supplementary materials S4-5). All five pre-contact samples, the soil sample, one post-contact sample putatively negative for *S. enterica* based on our MALT screening, and both rich UDG-treated (DNA damage removed) and non-UDG treated libraries from the three samples considered positive for *S. enterica* (Tepos_10, Tepos_14, Tepos_35) were included in the capture (Supplementary materials S5). Negative controls were pooled and captured on a separate array. The captured products for the samples and negative controls were paired-end sequenced using 2x75bp cycles on Illumina platforms to a depth of 19,758,038-192,213,420 reads per sample and 670,824-3,859,834 reads per negative control (tables S5).

Mapping and genotyping of the captured and sequenced reads was performed using the *S.* Paratyphi C genome reference (NC_012125.1). Capture of *S. enterica* DNA was successful for the three positive samples, but all other samples and negative controls were considered to be absent of *S. enterica* DNA (Supplementary materials S6; table S5). Genomes Tepos_10, Tepos_14 and Tepos_35, respectively, covered 83%, 97% and 98% of the reference at a minimum of 5-fold coverage and had an average coverage of 22-, 27- and 77-fold. Artificial reads generated *in silico* for 23 complete genomes included in the probe design were also mapped to the *S.* Paratyphi C RKS4594 reference (table S4) and phylogenetic comparison revealed that the three ancient genomes clustered with *S.* Paratyphi C (Supplementary materials S6; fig. S2). The phylogenetic positioning was retained when the whole dataset was mapped to and genotyped against the *S.* Typhi CT18 reference genome (NC_003198.1) (fig. S3; table S6), which is the most common cause of enteric fever in humans today. This result excludes the possibility of a reference bias. Oddly, despite all three samples being contemporaneous, the Tepos_10 genome was observed to contain many more derived positions than the other two ancient genomes. An investigation of heterozygous variant calls showed that Tepos_10 has a much higher number of heterozygous sites than Tepos_14 and Tepos_35. We believe this is best explained by the presence of genetically similar non-target DNA that was co-enriched in the capture for this sample alone. Based on the pattern of allele frequencies, this genome was excluded from downstream analyses (Supplementary materials S8; fig. S4). Subsequent phylogenetic tree construction with 1000 bootstrap replicates revealed 100% support and branch shortening for the Tepos_14 and Tepos_35 genomes in all phylogenies, supporting their ancient origin (Figs. 2, S5, S6).

SNP (single nucleotide polymorphism) analysis for the ancient strains was carried out in comparison to our reference dataset (table S4). The complete dataset consisted of 203,257 variant positions. Our analyses identified 681 positions present in one or both of the ancient genomes, where 131 are unique to the ancient lineages (Supplementary materials S9). Furthermore, 128 of these unique SNPs are shared between Tepos_14 and Tepos_35, thus indicating their close relationship and shared ancestry. The *ydiD* gene involved in the breakdown of fatty acids (*34*) and the *tsr* gene related to the chemotaxic response system (*35*) were found to contain multiple non-synonymous SNPs (nsSNPs) unique to the ancient genomes within the dataset (Supplementary materials S9). We detected six homoplastic and four tri-allelic variant positions in the ancient genomes in comparison to the modern genome dataset (Supplementary materials S9; tables S9A-B).

Analysis of insertions and deletions (indels) revealed regions in the ancient genomes that are comparatively absent in the modern *S.* Paratyphi C RKS4594 genome (Supplementary materials S10; table S10). One such region consists of five genes, *pilS*, *pilU*, *pilT*, *pilV* and *rci*. These genes form part of the *pil* operon located in Salmonella Pathogenicity Island 7 (SPI-7). In *S.* Typhi the Rci recombinase acts upon two 19bp inverted repeats in the *pilV* gene causing a rapid shuffling between two PilV protein states. This influences formation of the type IVB pili, causing them to be either malformed or not synthesized (*36*), thus contributing to bacterial self-aggregation, a phenomenon thought to aid in the invasion of host tissues (*37*). Our ancient genomes have the intact 19bp inverted repeats. Most modern S. Paratyphi C strains have 20bp inverted repeats that cause a mutational locking of *rci*’s shuffling function. For some strains, such as the *S*. Paratyphi C RKS4594 reference, these genes are either absent or degenerate (*37, 38*) (Supplementary materials S10). Additionally, analysis of *S. enterica* effector protein genes revealed that the *pipB2* gene appears to be fully present in the ancient strains, where it is covered 92% and 94% for Tepos_14 and Tepos_35 respectively, compared to the *S.* Paratyphi C RKS4594 genome where only 77% of *pipB2* is covered (Supplementary materials S11; figure S7; table S11).

The *S.* Paratyphi C RKS4594 strain harbors a virulence plasmid, pSPCV, which was included in our capture design. It is present at 56- and 178-fold average coverage for Tepos_14 and Tepos_35, respectively (Supplementary materials S12; table S12). pSPCV bears high genetic similarity to virulence plasmids present in the *S. enterica* serovars *S.* Choleraesuis (pSCV50 and pKDSC50) and *S.* Typhimurium (pSLT) (*39*). Comparison of filtered SNPs between the two ancient and four modern plasmids revealed two nsSNPs unique to the ancient plasmids (Supplementary materials S12; table S13).

Interpretations of ethnohistorical and archaeological evidence have suggested hemorrhagic fever, some form of typhus or typhoid fever (from the Spanish “*tifus mortal”*), measles, and bubonic plague as potential causes of the *cocoliztli* epidemic of 1545-50 CE (*10, 12, 28, 30, 40*). Previous investigation of sequencing data generated from the Teposcolula-Yucundaa material did not identify DNA traces of ancient pathogens; however, *S. enterica* was not considered as a candidate (*41*). Here we have reconstructed two high coverage genomes of ancient *S.* Paratyphi C from individuals of indigenous origin (Tepos_14 and Tepos_35) buried in the *Grand Plaza* epidemic cemetery at Teposcolula-Yucundaa. This indicates that *S.* Paratyphi C (enteric fever) was circulating in the indigenous population during the *cocoliztli* epidemic of 1545-50 CE. As demonstrated here, MALT offers a sensitive approach for screening non-enriched sequence data in search for unknown candidate bacterial pathogens involved in past disease outbreaks, even to the exclusion of a dominant environmental microbial background. Most importantly, it offers the advantage of extensive genome-level screening without the need to specify a target organism (*26*), thus avoiding ascertainment biases common to other screening approaches. Our exclusive focus on bacterial taxa, however, limits our resolution in identifying other infectious agents that may have acted either independently or synergistically with *S.* Paratyphi C during the Teposcolula-Yucundaa epidemic.

We confidently exclude an environmental organism as the source for our ancient genomes on the basis that 1) *S.* Paratyphi C is highly human-specific, 2) it is not known to freely inhabit soil (and additionally our soil sample was negative for *Salmonella* both during screening and after capture), 3) the deamination patterns observed for the ancient human and *S.* Paratyphi C reads are characteristic of authentic ancient DNA, and 4) the ancient *S.* Paratyphi C genomes display expected branch-shortening in all constructed phylogenies. Moreover, we recovered both ancient genomes from the pulp-chambers of teeth collected *in situ*, thus increasing the likelihood of our having identified a bacterium that was circulating in the blood of the victim at the time of death. Post-burial disturbance was limited by the graves of the *Grand Plaza* having been dug directly into the thickly paved floor at the site (*27, 28*), which is itself located on a mountain ridge above all other settlement and agricultural sites. Furthermore, historical records indicate that Teposcolula-Yucundaa was abandoned shortly after the epidemic ended, ca. 1552 (*27, 28*).

*S.* Paratyphi C is one of over 2600 identified *S. enterica* serovars distinguished by their antigenic formula (*42*). Only a few of these serovars are highly restricted to the human host (*42*) and consist of *S.* Typhi and *S.* Paratyphi A, B, C, all of which cause enteric fever. Today *S.* Typhi and *S.* Paratyphi A cause the majority of reported cases (*43*) while *S.* Paratyphi C is rarely reported (*39*). Infected individuals continue to shed bacteria long after the termination of symptoms (*42*), and in cases of infection with *S.* Typhi, as many as 1-6% become asymptomatic carriers (*44*) with similar percentages proposed for *S.* Paratyphi (*42*), a characteristic that is rare amongst human infectious diseases. Following the hypothesis that this disease was introduced via European contact, it is conceivable that asymptomatic European carriers who withstood the cross-Atlantic voyage could have introduced *S.* Paratyphi C to Mesoamerican populations in the 16^th^ century. First hand descriptions of the 1545 *cocoliztli* epidemic suggest that both European and Mixtec individuals were susceptible to the disease (*12, 45*), with estimates of 60-90% decline for the Mixtec population (*12*).

The additional SPI-7 genes detected through indel analysis are reported to vary in presence/absence amongst modern *S.* Paratyphi C strains (*37, 39*), and are suspected to cause increased virulence when the inverted repeats in *pilV* allow the Rci recombinase to shuffle between its two protein states (Supplementary materials S10). This may support an increased capacity for our ancient strains to cause an epidemic outbreak. However, *S.* Paratyphi A, which lacks the entire SPI-7 region, is currently one of the main causes of enteric fever worldwide (*39, 46*), and the overall mechanisms through which *S.* Paratyphi causes enteric fever remain unclear. The nsSNPs in the *ydiD* and *tsr* genes may signify adaptive processes, and comparison with a greater number of *S.* Paratyphi C genomes may help to clarify this (*47*).

Although a local origin for the *cocoliztli* disease has been proposed elsewhere (*10, 48*), we believe the *S.* Paratyphi C strains identified here were likely introduced from the Old World during the contact era, as evidenced by the existence of this human pathogen in Europe in 1200 CE (*47*). Historical accounts offer little perspective on its origin since neither the indigenous population nor the European colonizers had a pre-existing name for the disease (*11, 12, 28*). The Spanish called it *pujamiento de sangre* (‘full bloodiness’), while the indigenous Aztec population of Central Mexico named it *cocoliztli*, meaning ‘pestilence’ in Nahuatl (*11*). The name *cocoliztli* was also later applied to a pandemic in 1576-1581, which was described first-hand by a Spanish survivor of both epidemics as being caused by the same disease (*45*).

Enteric fever was first determined to be distinct from typhus in the mid-nineteenth century, hence little is known about the prior severity and worldwide incidence of enteric fever (*49*). Today, outbreaks predominantly occur in developing countries, where mortality rates from *S.* Typhi are reported to have reached as high as 30-50% (*50*). *S.* Typhi and *S.* Paratyphi are commonly transmitted through the fecal-oral route via ingestion of contaminated food or water (*51*). Changes imposed under Spanish rule such as forced relocations under the policy of *congregación*, altered living arrangements, and new subsistence farming practices (*27, 28*) compounded by drought conditions (*52*) could have disrupted existing hygiene measures, thus facilitating *S.* Paratyphi C transmission.

Our study represents a first step towards a molecular understanding of disease exchange in contact era Mexico. The 1545 *cocoliztli* epidemic is regarded as one of the most devastating epidemics in New World history (*12, 52*). Our findings contribute to the debate concerning the causative agent of this epidemic at Teposcolula-Yucundaa, where we propose that *S.* Paratyphi C be considered. Our novel use of MALT to identify ancient *Salmonella* DNA within a complex background of environmental microbial contaminants speaks to the power of this approach, which will only grow as the number of publicly available reference genomes continues to expand. This method can also be used to screen for a wider array of human pathogens such as DNA viruses and eukaryotes and may be eminently useful for studies wishing to identify pathogenic agents involved in ancient and modern disease, particularly in cases where candidate organisms are not known *a priori*

## Acknowledgments

This work was supported by the Max Planck Society (J.K.), the European Research Council (ERC) starting grant APGREID (to J.K.), Social Sciences and Humanities Research Council of Canada postdoctoral fellowship grant 756-2011-501 (to K.I.B.), and the Mäxi Foundation (M.G.C.). We thank the The Archaeology Council at Mexico’s National Institute of Anthropology and History (INAH) and the Teposcolula-Yucundaa Archaeological Project for sampling permissions. We are grateful to Antje Wissgott, Guido Brandt and Verena Schuenemann for assistance with laboratory work, Annette Günzel for providing graphic support, and Rodrigo Barquera for thoughts and discussion on the manuscript. Part of the data storage and analysis was performed on the computational resource bwGRiD Cluster Tübingen funded by the Ministry of Science, Research and the Arts Baden-Württemberg, and the Universities of the State of Baden-Württemberg, Germany, within the framework program bwHPC. K.I.B, M.G.C., A.H., N.T. and J.K. conceived the investigation. K.I.B, A.H. Å.J.V., M.G.C. and J.K. designed experiments. N.M.R.G. provided archaeological information and drawings, submitted INAH permits and assisted in the sampling processes. Å.J.V., M.G.C., M.A.S. and K.I.B. performed laboratory work. Å.J.V., A.H., K.I.B., C.W. and A.A.V. performed analyses. Å.J.V. and K.I.B. wrote the manuscript with contributions from all authors. Data was submitted to the European Nucleotide Archive under accession no. [XXXX]. The authors declare no competing interests.

**Table 1.** Overview of captured sample libraries from the Grand Plaza (Contact) and churchyard (Pre-contact), and mapping statistics.

| Sample ID | Burial Site | Library treatment | # Merged Paired-End Reads | # Unique Mapping Reads | Endogenous DNA (%) - quality filtered reads | Mean Fold Coverage | % of Genome Covered at least 5-fold |
| --- | --- | --- | --- | --- | --- | --- | --- |
| **Tepos_10** | Grand Plaza | non-UDG | 14,067,111 | 289,468 | 20.2 | 3.1 | 21.3 |
|  |  | UDG | 58,806,841 | 2,056,326 | 16.1 | 22.6 | 83.6 |
| **Tepos_14** | Grand Plaza | non-UDG | 16,958,495 | 746,869 | 23.1 | 8.4 | 76.7 |
|  |  | UDG | 62,441,891 | 2,474,496 | 18.4 | 25.8 | 96.8 |
| **Tepos_35** | Grand Plaza | non-UDG | 24,518,209 | 1,430,852 | 31.6 | 19.2 | 95.8 |
|  |  | UDG | 81,358,355 | 5,852,171 | 30.3 | 76.6 | 97.9 |
| **Tepos_32** | Churchyard | non-UDG | 20,726,258 | 1,468 | 0.1 | 0.0 | 0.1 |
| **Tepos_43** | Churchyard | non-UDG | 25,565,135 | 1,318 | 0.0 | 0.0 | 0.1 |
| **Tepos_45** | Churchyard | non-UDG | 14,373,878 | 1,864 | 0.1 | 0.0 | 0.1 |
| **Tepos_48** | Churchyard | non-UDG | 14,204,209 | 1,140 | 0.1 | 0.0 | 0.1 |
| **Tepos_57** | Churchyard | non-UDG | 19,961,358 | 1,598 | 0.1 | 0.0 | 0.1 |
| **Tepos_27** | Grand Plaza | non-UDG | 14,115,775 | 1,314 | 0.1 | 0.0 | 0.1 |
| **soil** | Grand Plaza & churchyard | non-UDG | 7,149,517 | 67 | 0.0 | 0.0 | 0.0 |

## Supplementary Materials

Materials and Methods

Figures S1-S7

Tables S1-S13

References (*53-83*)

